# Optotracers to study biofilms in host-relevant systems and non-conventional treatments using *Staphylococcus aureus* wound biofilms as a case study

**DOI:** 10.1101/2023.02.02.526914

**Authors:** Nijamuddin Shaikh, Karishma S Kaushik

## Abstract

Optotracers have a wide-range of applications in the detection, visualization, and characterization of biofilms. More recently, optotracers have been used in antibiotic susceptibility assays for biofilms and for the detection of biomarkers in clinical biofilm infections. This is particularly important, given that the field is increasingly focused on the evaluation of novel anti-biofilm agents and the study of biofilms in host-relevant systems. Given this, the possibility of using optotracers for an expanded set of biofilm focus areas is important to explore. In this study, we examine the application of the optotracer EbbaBiolight 680 to study biofilms in a host-relevant system and for the evaluation of a non-conventional anti-biofilm remedy. Using *Staphylococcus aureus* wound biofilms as a case study, we leverage a previously built *in vitro* 4-D wound microenvironment platform and a plant-based wound remedy. We find that EbbaBiolight 680 can be used to visualize *S. aureus* biofilms, likely detecting both bacterial cells and bacterial EPS components. Further, the optotracer can be used to evaluate and quantify the effects of the plant-based wound remedy on *S. aureus* biomass formation. However, in the 4-D wound microenvironment, EbbaBiolight 680 detected host cellular and matrix elements, which confounded the detection of biofilms. Taken together, this study opens the possibility of using optotracers as screening tools for the identification of novel anti-biofilm treatments and underscores the need for modifications for their use in host-relevant systems.

## Introduction

Optotracers are fluorescent visualization molecules that can be used for the real-time detection, visualization, characterization, and quantification of biofilms [1–3]. When added to biofilm growth conditions or assays, optotracers have been reported to detect both bacterial cells and biofilm extracellular polymeric substance (EPS) components, which enables the *in situ* visualization of biofilm formation, structure, and development [4–6]. Given this ability to visualize live biofilms in a non-disruptive manner, optotracers have been used to study biofilm formation and features in a range of pathogens, including *Salmonella*, Uropathogenic *Escherichia coli* (UPEC) and *Staphylococcus aureus* [1,2,4,6,7]. In addition to detection and visualization of biofilms, the use of optotracers has expanded to applications in biofilm antibiotic susceptibility assays and biofilm markers in clinical diagnostics [1,4,7]. Notably, this raises the possibility of using optotracers in the evaluation of novel anti-biofilm agents as well as in the study of biofilms under conditions that mimic the clinical infection state. This is particularly important in the broader context of the biofilm field, which is increasingly focused on studying biofilms under host-relevant conditions and identifying non-conventional anti-biofilm treatments [8–16].

In this study, we explored the application of the optotracer EbbaBiolight 680 in the study of biofilms in an *in vitro* host-relevant system and for the evaluation of a potential non-conventional anti-biofilm remedy using *Staphylococcus aureus* biofilms in wounds as a case study [17]. EbbaBiolight 680 is a red/far-red optotracer molecule known to bind to repetitive motifs in bacterial cell and EPS components, following which it produces a conformation-dependent fluorescence emission [18]. We observed that EbbaBiolight 680 could detect and visualize the bacterial cells and bacterial EPS components of *S. aureus* biofilms. To examine the role of EbbaBiolight 680 in identifying potential non-conventional anti-biofilm treatments, we evaluated a previously reconstituted and characterized plant-based wound remedy of *Ocimum tenuiflorum* and sesame oil (OTE-SO) [19]. We observed that EbbaBiolight 680 could be used to visualize and quantify effects of the OTE-SO wound remedy on *S. aureus* biomass and biofilm structure, with differential effects on two different *S. aureus* strains. To examine the use of EbbaBiolight 680 for biofilms in host-relevant systems, we leveraged our previous *in vitro* 4-D wound microenvironment platform, consisting of co-cultured host cells (fibroblasts and keratinocytes) and an *in vitro* wound milieu (IVWM) [12,20]. For this, *S. aureus* biofilms (three different strains) were grown in the 4-D wound microenvironment, and EbbaBiolight 680 was used to visualize the biofilms and study the effects of the antibiotic vancomycin, often used in the treatment of *S. aureus* biofilms [21,22]. We observed that EbbaBiolight 680 also detected the host cellular and matrix elements in the system, seen as visualization of fibroblasts and keratinocytes, and components of the IVWM, which confounded the visualization of the biofilms and assessment of the effects of the antibiotic.

## Materials and Methods

### Bacterial strains and growth conditions

The *S. aureus* strain AH13 (pAH13::GFPuvr, ErmR) was a gift from Dr. Derek Fleming and Dr. Kendra Rumbaugh (Texas Tech University Health Science Center, Lubbock, Texas, USA) [23]. *S. aureus* MCC 2043 was purchased from National Centre for Microbial Resource (NCMR), Pune, India. *S. aureus* S-235 was a gift from Dr. Joey Shepherd (School of Clinical Dentistry, University of Sheffield, UK) [24]. For all experiments, *S. aureus* strains were grown in Luria-Bertani (LB) agar (SRL, India, 474236) or buffered Tryptic Soy Broth (bTSB) [10 g/ml K_2_HPO_4_ (Himedia, India, GRM168), 1 g/L KH_2_PO_4_ (Himedia, India, MB050) in TSB (Himedia, India, M1263)] and incubated at 37°C for 18-20 hours under static conditions for the agar plates and shaking conditions for the broths [10 μg/mL erythromycin (Himedia, India, TC024) was added for *S. aureus* AH13].

### Host cell culture and maintenance

Primary human dermal fibroblasts (HDFa) were purchased from PromoCell (Germany, C-12302) and cultured in Fibroblast Growth Medium (FGM) containing 2% fetal bovine serum (FBS) (Cell Applications, USA 116– 500). The immortalized epidermal keratinocyte cell line (HaCaT) was a gift from Dr. Madhur Motwani (Linq Labs, Pune, India) and was cultured in Keratinocyte serum-free Growth Medium (KGM) (Cell Applications, USA, 131–500A) supplemented with 1% FBS (Gibco, Brazil, 10270106). Cells were grown in tissue-culture treated, cell culture flasks (Tarsons, Korea, 950040) at 37°C in a 5% CO_2_ humidified incubator.

### Visualizing *S. aureus* biofilms with EbbaBiolight 680

Optical densities of overnight cultures of *S. aureus* AH13 (grown with 10 μg/ml erythromycin) and *S. aureus* MCC 2043 were measured at 600 nm. Each of the bacterial suspensions were diluted in antibiotic-free bTSB to obtain a density of 1×10^7^ cells/ml. Of the diluted bacterial suspensions, 100 μL (~10^6^ cells) were added to individual wells of a 96-well flat, clear bottom, tissue-culture treated, black polystyrene microplate (Corning, USA, 3603) and incubated at 37°C in a static incubator. After one hour of incubation, the supernatant medium was gently removed and replaced with bTSB containing a 1:1000 dilution of EbbaBiolight 680 (test sample from Ebba Biotech AB, Sweden, E680) and incubated at 37°C in a static incubator for 24 hours. Further, bTSB medium with *S. aureus* MCC 2043 (without EbbaBiolight 680) and with EbbaBiolight 680 only (without bacteria) were also set up.

### Preparation of the OTE-SO remedy

The OTE-SO remedy was reconstituted as previously described, with certain modifications [19]. Briefly, the leaves of *Ocimum tenuiflorum* were made into a paste, and mixed with distilled water and sesame oil (Prabhat teel oil, Hanuman Trading Company, India) in a ratio of 1:16:4 (by volume). The mixture was boiled till the water completely evaporated, the residual extract-containing oil was filtered and stored at room temperature. The reconstituted OTE-SO remedy was mixed with absolute ethanol (Changshu Hongsheng Fine Chemical Co. Ltd.) and dimethyl sulphoxide (DMSO) (Thermo Scientific, France, 85190) in a ratio of 2:1:1 (by volume) and incubated at 37°C for 24 hours under shaking conditions. The mixture was subsequently stored at room temperature for 24 hours to attain phase separation. After 24 hours, the upper (oil) phase was discarded and the residual mixed solution was used as the OTE-SO remedy.

### Evaluation of OTE-SO remedy against *S. aureus* biofilm formation with EbbaBiolight 680

As previously described, ~10^6^ cells of *S. aureus* MCC 2043 or *S. aureus* AH13 (GFP) diluted in antibiotic-free bTSB were added to individual wells of 96-well, flat, clear bottom, tissue-culture treated, black polystyrene microplate and incubated at 37°C in a static incubator. After 1 hour of incubation, the supernatant medium was carefully removed and replaced with bTSB containing 1:1000 dilution of EbbaBiolight 680 and 1% of the OTE-SO remedy (prepared as above and diluted in bTSB) and incubated at 37°C in a static incubator for 24 hours.

### Assembly of the 4-D wound microenvironment

The 4-D wound microenvironment was assembled with co-cultured host cells (HDFa+HaCaT) and an IVWM as previously described [12]. Briefly, nearly equal counts of HDFa and HaCaT cells (~5000 cells each) were added to a single well of a 96-well, transparent bottom, tissue culture treated, black microplate (Corning, USA, 3603) and grown at 37°C in a 5% CO_2_ humidified incubator for 72 hours [25]. Following this, the confluent cells were fixed in 4% paraformaldehyde (PFA) (Sigma, USA, 158127) and stained with 4’,6-Diamidino-2-Phenylindole, Dihydrochloride (DAPI) (Invitrogen, Germany, D1306). The IVWM was reconstituted as previously described, with FBS as the base components and the addition of host matrix factors (collagen, fibronectin, fibrinogen) and biochemical factors (lactic acid and lactoferrin) [12,20]. Reconstituted IVWM was freshly prepared prior to use.

### Growth of *S. aureus* biofilms in the 4-D wound microenvironment with EbbaBiolight 680

Bacterial suspension of ~10^6^ cells of *S. aureus* MCC 2043, *S. aureus* AH13 (GFP), *S. aureus* S-235 in IVWM were each added to different fixed co-cultured (HDFa+HaCaT) host cell scaffolds and incubated at 37°C in a static incubator for one hour. Subsequently, the supernatant medium from each well was gently removed and replaced with 100 μL of freshly-prepared IVWM containing 1:1000 dilution of EbbaBiolight 680. This 4-D wound microenvironment system, consisting of fixed host cells, IVWM, and *S. aureus*, was incubated at 37°C in a static incubator for 24 hours.

### Evaluation of vancomycin against *S. aureus* biofilms in the 4-D wound microenvironment with EbbaBiolight 680

As described, ~10^6^ cells of *S. aureus* AH13, *S. aureus* MCC 2043, and *S. aureus* S-235 in IVWM were each added to fixed co-cultured (HDFa+HaCaT) host cell scaffolds and incubated at 37°C under static conditions. After one hour, the supernatant medium was carefully replaced with IVWM containing a 1:1000 dilution of EbbaBiolight 680 and 100 μg/mL vancomycin (from a stock concentration of 10 mg/mL). The entire set-up was incubated at 37°C under static conditions for 24 hours.

### Evaluating EbbaBiolight 680 in the presence of host cellular and matrix factors (in the absence of *S. aureus* biofilms)

To evaluate EbbaBiolight 680 in the presence of cellular and matrix factors, a 1:1000 dilution of the optotracer in freshly-prepared IVWM was added to the fixed host cell scaffolds (no bacterial inoculum was added) and the entire system was incubated at 37°C in a static incubator for 24 hours. To evaluate EbbaBiolight 680 in the presence of the host cell scaffolds alone, a 1:1000 dilution of the optotracer in bTSB was added to the fixed host cell scaffolds, the set-up was incubated at 37°C under static conditions for 24 hours.

### Confocal microscopy and image acquisition

At 24 hours, *S. aureus* biomass across different conditions (as described above) was visualized using confocal laser scanning microscopy (Leica, Germany, LASX TCS SP). For each well, the imaging was done at the approximate center. To visualize EbbaBiolight 680, a 561 nm excitation filter and 586-656 nm emission filter was used. To visualize GFP-tagged *S. aureus* AH13 an excitation filter of 488 nm and emission filter of 497-542 nm was used. To visualize DAPI-stained host cells, a 405 nm excitation filter and 414-485 nm emission filter was used. The biomass was imaged using Z-stacks starting at the lower end of the biomass and up to of a total of 100 μm thickness, using a Z-step size of 5 μm.

### Image processing and analysis

Image analysis was done using the open-source image analysis tool, ImageJ [26]. For this, raw LIF files for each *S. aureus* strain and study condition were imported into ImageJ. In ImageJ, maximum intensity Z-projections were obtained, which were subject to thresholding by the Isodata method [27]. To calculate biomass, the Mean Gray Value (MGVs) calculations in ImageJ were used [12]. The MGV of the red channel was used to calculate the biomass obtained by EbbaBiolight 680 emission and the MGV of the green channel was measured to calculate the biomass due to GFP fluorescence.

### Statistical analysis

Statistical analysis was performed using Microsoft Office 2021 (MS Excel) for Windows [28]. A single-factor ANOVA test was performed and a p-value of ≤ 0.05 was considered significant.

## Results and Discussion

### The optotracer EbbaBiolight 680 detects bacterial cells, and possibly EPS components, of *S. aureus* biofilms

The red/far-red optotracer EbbaBiolight 680 was added to bTSB medium with *S. aureus* MCC 2043 or GFP-tagged *S. aureus* AH13, and biofilms were visualized after 24 hours of growth. As seen in **Figure 1A**, *S. aureus* MCC 2043 biomass was visualized as red fluorescence, indicating the binding of EbbaBiolight 680 to components in the biomass. Further, the biofilms appeared as heterogeneous clumps or aggregates of biomass. For GFP-tagged *S. aureus* AH13, EbbaBiolight 680 was observed to detect large clumps of biomass **(Figure 1B)**. Notably, in comparison with the green channel (for detection of GFP-tagged bacteria), the red fluorescence appears dense and dispersed across regions without bacterial cells, indicating the possibility of EbbaBiolight 680 binding to bacterial EPS components in *S. aureus* biomass. In bTSB medium with only *S. aureus* MCC 2043 (no optotracer), no red fluorescence emission was observed **(Figure 1C)**. Further, minimal red fluorescence emission was observed when EbbaBiolight 680 was added to bTSB medium without *S. aureus* (no bacterial inoculum) **(Figure 1D)**. Together, this indicates that the EbbaBiolight 680 optotracer detects bacterial cells, and possibly bacterial EPS components, in biofilms formed by two different strains of *S. aureus*. The optotracer EbbaBiolight 680 has been reported to bind to EPS components, such as the amyloid protein curli, in *Salmonella* biofilms [18]. In *S. aureus* biofilms, the binding targets of EbbaBiolight 680 could include a range of EPS components; in addition, the binding targets of EbbaBiolight 680 in Gram-positive bacterial cells, namely cell wall components, could also be shed into the EPS [29,30]. Further characterizing the presence and identity of the specific binding targets in the EPS could lead to a range of applications of EbbaBiolight 680 in the study of *S. aureus* biofilms.

**Figure 1:**
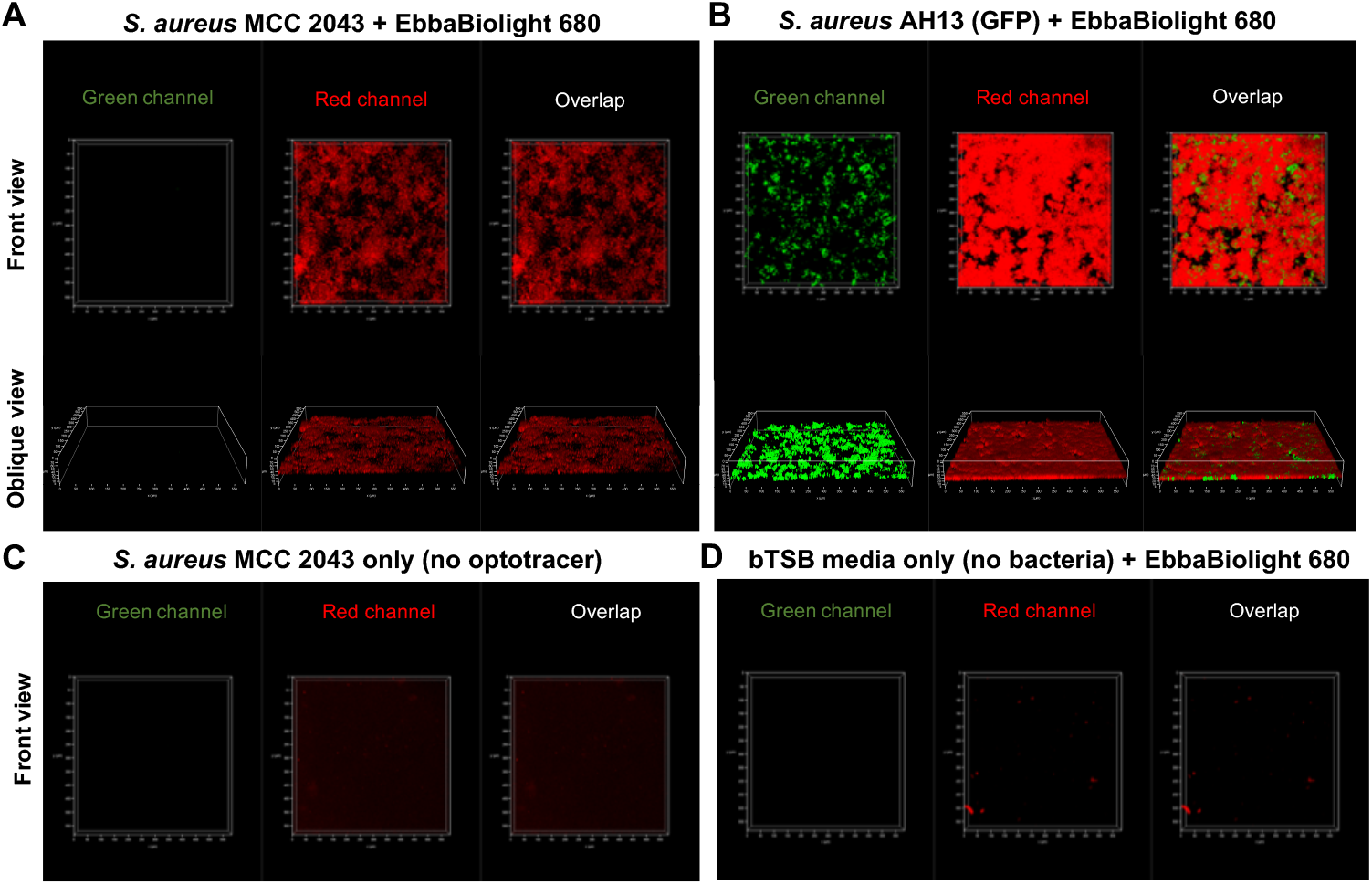
The optotracer EbbaBiolight 680 detects bacterial cells, and possibly EPS components, of *S. aureus* biofilms. The red/far-red optotracer EbbaBiolight 680 was added to bTSB medium with *S. aureus* MCC 2043 or GFP-tagged *S. aureus* AH13, and biofilms were visualized after 24 hours of growth. **(A)** EbbaBiolight 680 detects *S. aureus* MCC 2043 biofilms, seen as clumps or aggregates with red fluorescence emission. **(B)** EbbaBiolight 680 detects biofilms formed by GFP-tagged *S. aureus* AH13, seen as large and dense clumps with red fluorescence emission. The red fluorescence emission is also seen in regions without bacterial cells (seen with GFP) and possibly indicates EbbaBiolight 680 detection of EPS components. **(C)** For *S. aureus* MCC 2043 biofilms, in the absence of EbbaBiolight 680, no red fluorescence is observed. **(D)** In bTSB medium alone without bacterial inoculum, addition of EbbaBiolight 680 does not produce red fluorescence.

### The optotracer EbbaBiolight 680 can be used to detect changes in *S. aureus* biomass and biofilm structure with a non-conventional wound remedy

To study the role of optotracers in the evaluation of non-conventional anti-biofilm treatments, the reconstituted plant-based remedy of *Ocimum tenuiflorum* and sesame oil (OTE-SO) was used [19]. Briefly, EbbaBiolight 680 was added to bTSB medium with *S. aureus* MCC 2043 or GFP-tagged *S. aureus* AH13, and after one hour the medium was replaced with the reconstituted OTE-SO remedy. The plant-based OTE-SO composition is based on historical medical prescriptions for non-healing wounds, and has been previously reconstituted and characterized for effects on *S. aureus* (GFP-tagged AH13) biofilm formation [19]. As seen in **Figure 2A**, in the presence of the OTE-SO remedy, *S. aureus* MCC 2043 formed less-dense biofilms (as compared with untreated biofilms in **Figure 1A**), with the biofilm aggregates seen to be smaller and more discrete. Further, when the biomass change is quantified (measured as MGV of the red channel in ImageJ), the OTE-SO treated biofilms show a significant decrease in biomass, as compared with the untreated biofilms (error bars represent SEM, n=3, technical replicates) **(Figure 2B)**. On the other hand, on treatment with OTE-SO, *S. aureus* AH13 (GFP) biofilms were observed in large clumps, similar to that under untreated conditions **(Figure 2C)**. Along these lines, biomass measured across the untreated **(Figure 1B)** and treated biofilms (measured as MGV of the red channel in ImageJ) also showed no significant difference (error bars represent SEM, n=3, technical replicates) **(Figure 2D)**. Further, the MGVs of the GFP intensities, indicative of the biomass of the bacterial cells alone, also showed no significant decrease in the presence of OTE-SO **(Figure 2E)**. While further evaluation of bacterial cell viability and CFU counts would be important, these observations indicate that EbbaBiolight 680 could be used to visualize and quantify effects of the non-conventional wound remedy OTE-SO on *S. aureus* biomass formation and biofilm structure, with differential effects across two different *S. aureus* strains. Further, the optotracer could enable the exploration of the effects of the treatments on both bacterial cells and bacterial EPS in biofilms. Together, this points to the possibility of using EbbaBiolight 680 as a screening tool for the semi-high throughput evaluation of novel and potential anti-biofilm agents, including non-conventional treatments.

**Figure 2:**
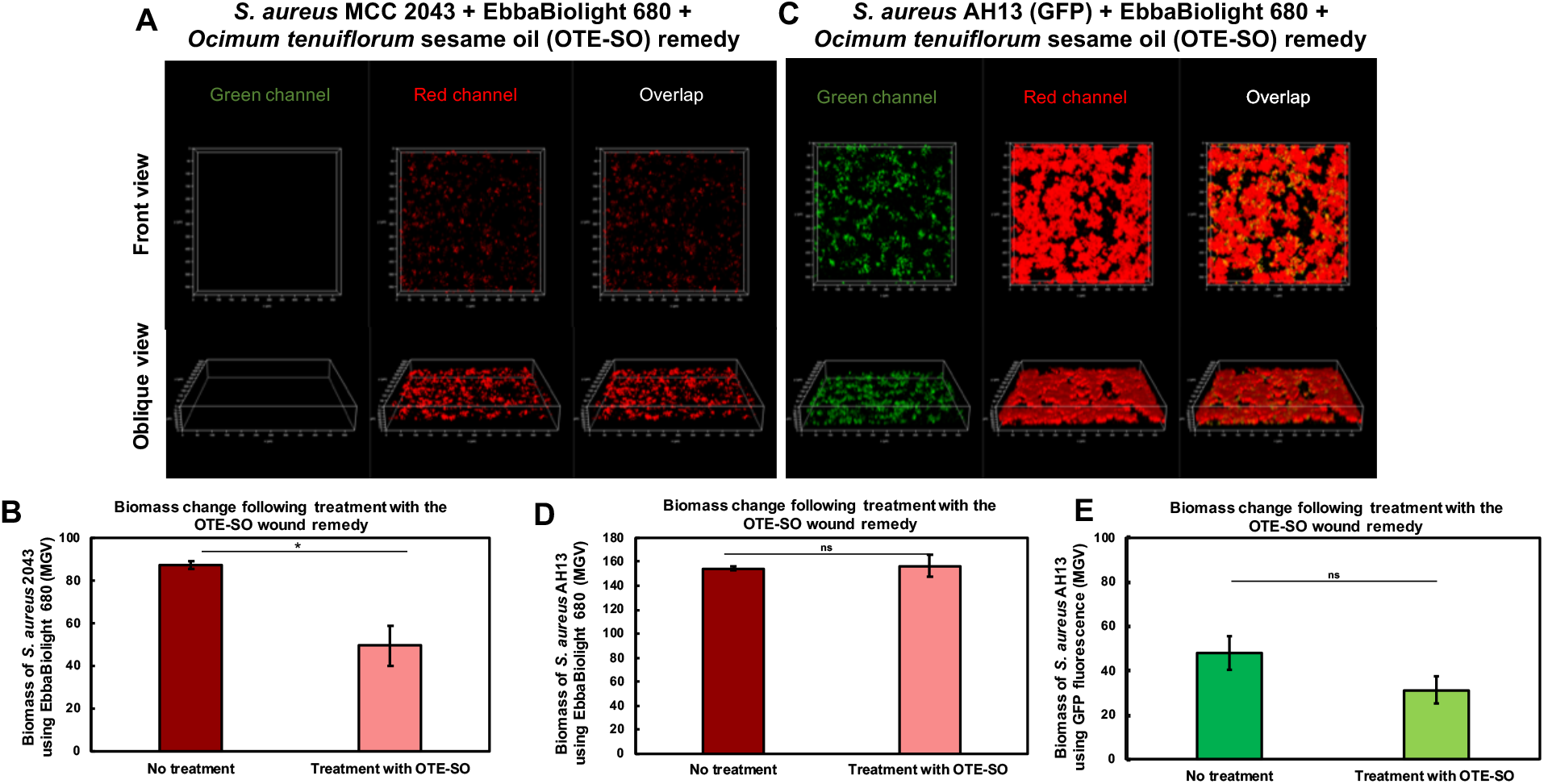
The optotracer EbbaBiolight 680 can be used to detect changes in *S. aureus* biomass and biofilm structure with a non-conventional wound remedy. The optotracer EbbaBiolight 680 was added *S. aureus* MCC 2043 or GFP-tagged *S. aureus* AH13 biofilms, and biofilm formation was examined in the presence of the reconstituted remedy of *Ocimum tenuiflorum* and sesame oil (OTE-SO). **(A)** *S. aureus* MCC 2043 treated with OTE-SO were observed to form less-dense biofilms, including smaller clumps or aggregates. **(B)** The effect of OTE-SO on *S. aureus* MCC 2043 biomass was quantified using MGV analysis (of the red channel) in ImageJ, and showed a reduction in biomass as compared with untreated biomass. **(C)** *S. aureus* AH13 (GFP) biofilms treated with OTE-SO were similar to that under untreated conditions, with large clumps of biomass. **(D)** The effect of OTE-SO on *S. aureus* AH13 (GFP) biofilms was quantified using MGV analysis (of the red channel) in ImageJ, and showed no significant difference in measurements as compared with untreated biomass. **(E)** For *S. aureus* AH13 (GFP) biofilms, the MGVs of GFP intensities showed no significant difference in the presence of OTE-SO. Error bars represent SEM, n=3, technical replicates; p-value of ≤ 0.05 was considered significant (*).

### In a host-relevant system, the optotracer EbbaBiolight 680 is unable to distinguish biofilms and assess the effects of antibiotic treatment

To examine the use of EbbaBiolight 680 for biofilms in host-relevant systems, we leveraged our previous *in vitro* 4-D wound microenvironment platform, consisting of cocultured host cells (HDFa+HaCaT) and an *in vitro* wound milieu (IVWM) [12,20]. As seen in **Figures 3A-C**, with *S. aureus* biofilms in the 4-D wound microenvironment, EbbaBiolight 680 produced red fluorescence emission across the entire system, resulting in the inability to distinguish the bacterial cells and biofilms. Notably, this was observed for three different strains, *S. aureus* MCC 2043, GFP-tagged *S. aureus* AH13, and the clinical strain *S. aureus* S-235. Further, when subject to treatment with the antibiotic vancomycin (100 μg/mL), the uniform red fluorescence emission across the microenvironment precluded the assessment of the effects of the antibiotic **(Figures 3D and 3E)**. This was observed for both the clinical strain *S. aureus* S-235 and GFP-tagged *S. aureus* AH13; notably, the MGV of the GFP fluorescence in *S. aureus* AH13 shows a decrease across the untreated and treated biofilms **(Figure 3F)**. Given this, in the 4-D wound microenvironment, EbbaBiolight 680 was unable to distinguish bacterial cells and biofilms, and in doing so, was unable to evaluate the effects of antibiotics in the system. Based on visual observations, the uniform red fluorescence emission across the 4-D wound microenvironment is likely due to optotracer detection and visualization of the non-biofilm components of the platform, which includes the IVWM and co-cultured host cells.

**Figure 3:**
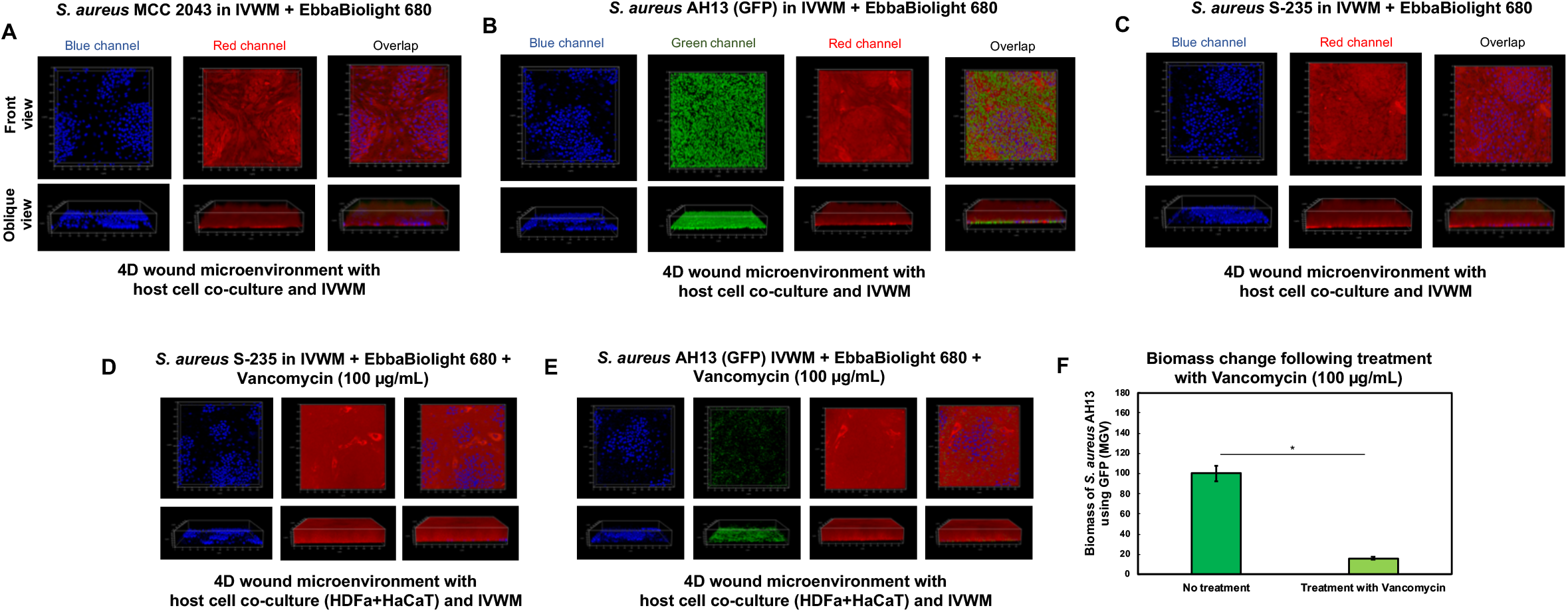
In a host-relevant system, the optotracer EbbaBiolight 680 is unable to distinguish biofilms and assess the effects of antibiotic treatment. To evaluate the use of EbbaBiolight 680 for biofilms in host-relevant systems, three different *S. aureus S. aureus* MCC 2043, GFP-tagged *S. aureus* AH13, and the clinical strain *S. aureus* S-235 were inoculated in the *in vitro* 4-D wound microenvironment platform, consisting of cocultured host cells (HDFa+HaCaT) and the IVWM. **(A-C)** For all three strains of *S. aureus* biofilms in the 4-D wound microenvironment, EbbaBiolight 680 produced red fluorescence emission across the entire system. **(D-E)** For *S. aureus* S-235 and GFP-tagged *S. aureus* AH13, a uniform red fluorescence of EbbaBiolight 680 was observed when biofilm formation was evaluated in the presence of the antibiotic vancomycin. **(F)**The MGVs of GFP fluorescence in *S. aureus* AH13 showed a decrease in biofilms treated with vancomycin, as compared with untreated biofilms. Error bars represent SEM, n=2, technical replicates; p-value of ≤ 0.05 was considered significant (*).

### The optotracer EbbaBiolight 680 detects host matrix and cellular elements, namely the IVWM and co-cultured host cells, in the 4-D wound microenvironment

To confirm the possibility of the optotracer detecting host matrix and cellular factors, EbbaBiolight 680 was added to the 4-D wound microenvironment system without *S. aureus* biofilms, as well as the co-cultured host cell system alone (without the IVWM and *S. aureus* biofilms). As seen in **Figure 4A**, in the presence of IVWM and the host cell scaffold, EbbaBiolight 680 produced a uniform red fluorescence across the 4-D wound microenvironment, likely indicating the detection of host factors in the IVWM. Further, in the absence of both the IVWM and *S. aureus* biofilms **(Figure 4B)**, EbbaBiolight 680 detected the co-cultured host cells, seen as red fluorescent HDFa and HaCaT cells with characteristic cellular structures and arrangements [25]. Taken together, EbbaBiolight 680 detected host cellular and matrix elements in the 4-D wound microenvironment, which confounded the optotracer visualization of biofilms and evaluation of antibiotic treatment in the host-relevant system. It is important to note that the IVWM contains FBS as the base component, along with matrix factors such as collagen, fibrinogen and fibronectin and biochemical factors, lactic acid and lactoferrin [20]. Further, dermal fibroblasts and epidermal keratinocytes are also known to secrete matrix proteins, including fibrinogen, fibronectin, collagen and keratin [31,32]. Finally, the EPS of *S. aureus* biofilms is known to consist of a range of proteins, including host matrix-binding proteins for fibrinogen and fibronectin [30]. Given this, it is possible that the repetitive motifs detected by EbbaBiolight 680 are also present in the structure of select components of the 4-D wound microenvironment [5,18,33,34].

**Figure 4:**
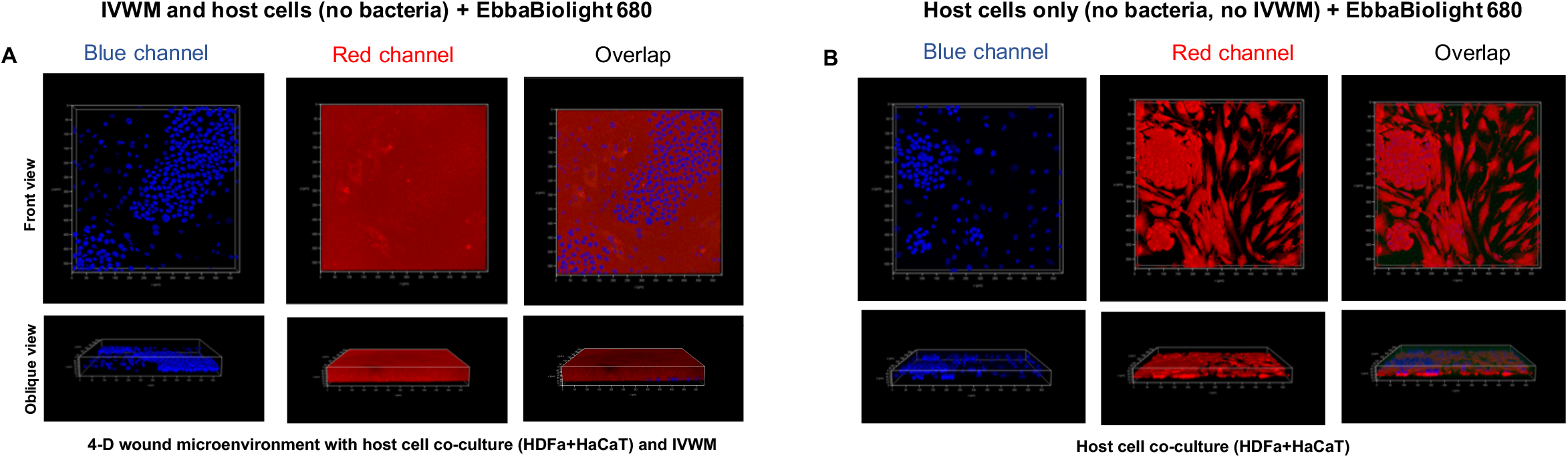
The optotracer EbbaBiolight 680 detects host matrix and cellular elements, namely the IVWM and co-cultured host cells, in the 4-D wound microenvironment. To confirm the possibility of EbbaBiolight 680 detecting host matrix and cellular factors, the optotracer was added to the 4-D wound microenvironment without *S. aureus* biofilms, as well as the co-cultured host cell system alone (without the IVWM and *S. aureus* biofilms). **(A)** In the absence of bacteria, EbbaBiolight 680 produced a uniform red fluorescence across the 4-D wound microenvironment. **(B)** In the presence of EbbaBiolight 680, the co-cultured host cells were seen as red fluorescent structures.

## Conclusions

The optotracer EbbaBiolight 680 can be used to visualize *S. aureus* biofilms, likely detecting both bacterial cells and bacterial EPS components. This enables the evaluation of novel and potential anti-biofilm agents, including non-conventional remedies, where it provides insights into the effects of these treatments on both bacteria and EPS elements. However, in host-relevant systems, the detection of host cellular and matrix factors confounds the detection of biofilms and evaluation of antibiotic treatments. Given the increasing relevance of studying biofilms under host-relevant conditions, optotracer molecules may need to be further modified prior to their use in these systems. These modified optotracers could also be applied to the study of biofilms directly from clinical material, including biopsies and exudates. We believe that this characterization of EbbaBiolight 680 using *S. aureus* wound biofilms as a case study will enable the application and adaptation of optotracers for a range of biofilm studies and biofilm-forming pathogens.

## Acknowledgements

We thank Smilla Huzell from Ebba Biotech AB, Sweden for the sample of EbbaBiolight 680 (optotracer), and the organizers of Eurobiofilms 2022, Palma de Mallorca, Spain for the opportunity to interact with the Ebba Biotech AB team. We thank Sujaya Ingale and M Vandana from NCL-Innovation Park, Pune, India for technical assistance with the confocal microscopy. We thank Dr. Anuradha Bandgar, Dr. Anuradha’s Ayurvedic Clinic, Pune, India for the OTE-SO whole remedies.

## Data availability statement

Data generated in this study will be made available by the authors upon request.

## References

[1] Eckert JA, Rosenberg M, Rhen M, Choong FX, Richter-Dahlfors A. An optotracerbased antibiotic susceptibility test specifically targeting the biofilm lifestyle of Salmonella. Biofilm 2022;4:100083. https://doi.org/10.1016/J.BIOFLM.2022.100083.

[2] Butina K, Tomac A, Choong FX, Shirani H, Nilsson KPR, Löffler S, et al. Optotracing for selective fluorescence-based detection, visualization and quantification of live S. aureus in real-time. Npj Biofilms and Microbiomes 2020 6:1 2020;6:1–12. https://doi.org/10.1038/s41522-020-00150-y.

[3] Parlak O, Richter-Dahlfors A. Bacterial Sensing and Biofilm Monitoring for Infection Diagnostics. Macromol Biosci 2020;20:2000129. https://doi.org/10.1002/MABI.202000129.

[4] Antypas H, Choong FX, Libberton B, Brauner A, Richter-Dahlfors A. Rapid diagnostic assay for detection of cellulose in urine as biomarker for biofilm-related urinary tract infections. Npj Biofilms and Microbiomes 2018 4:1 2018;4:1–7. https://doi.org/10.1038/s41522-018-0069-y.

[5] Choong FX, Bäck M, Fahlén S, Johansson LBG, Melican K, Rhen M, et al. Realtime optotracing of curli and cellulose in live Salmonella biofilms using luminescent oligothiophenes. Npj Biofilms and Microbiomes 2016 2:1 2016;2:1–11. https://doi.org/10.1038/npjbiofilms.2016.24.

[6] Butina K, Lantz L, Choong FX, Tomac A, Shirani H, Löffler S, et al. Structural Properties Dictating Selective Optotracer Detection of Staphylococcus aureus. ChemBioChem 2022;23:e202100684. https://doi.org/10.1002/CBIC.202100684.

[7] Rodríguez-Rojas A, Baeder DY, Johnston P, Regoes RR, Rolff J. Bacteria primed by antimicrobial peptides develop tolerance and persist. PLoS Pathog 2021;17. https://doi.org/10.1371/JOURNAL.PPAT.1009443.

[8] Bjarnsholt T, Whiteley M, Rumbaugh KP, Stewart PS, Jensen P, Frimodt-Møller N. The importance of understanding the infectious microenvironment. Lancet Infect Dis 2022;22:e88–92. https://doi.org/10.1016/S1473-3099(21)00122-5.

[9] Verderosa AD, Totsika M, Fairfull-Smith KE. Bacterial Biofilm Eradication Agents: A Current Review. Front Chem 2019;7:824. https://doi.org/10.3389/FCHEM.2019.00824/BIBTEX.

[10] Wu H, Moser C, Wang HZ, Høiby N, Song ZJ. Strategies for combating bacterial biofilm infections. International Journal of Oral Science 2014 7:1 2014;7:1–7. https://doi.org/10.1038/ijos.2014.65.

[11] Kadam S, Shai S, Shahane A, Kaushik KS. Recent Advances in Non-Conventional Antimicrobial Approaches for Chronic Wound Biofilms: Have We Found the ‘Chink in the Armor’? Biomedicines 2019, Vol 7, Page 35 2019;7:35. https://doi.org/10.3390/BIOMEDICINES7020035.

[12] Dhekane R, Mhade S, Kaushik KS. Adding a new dimension: Multi-level structure and organization of mixed-species Pseudomonas aeruginosa and Staphylococcus aureus biofilms in a 4-D wound microenvironment. Biofilm 2022;4:100087. https://doi.org/10.1016/J.BIOFLM.2022.100087.

[13] Lu L, Hu W, Tian Z, Yuan D, Yi G, Zhou Y, et al. Developing natural products as potential anti-biofilm agents. Chinese Medicine (United Kingdom) 2019;14:1–17. https://doi.org/10.1186/S13020-019-0232-2/TABLES/3.

[14] Rumbaugh KP, Ahmad I, editors. Antibiofilm Agents 2014;8. https://doi.org/10.1007/978-3-642-53833-9.

[15] Mhade S, Panse S, Tendulkar G, Awate R, Narasimhan Y, Kadam S, et al. AMPing Up the Search: A Structural and Functional Repository of Antimicrobial Peptides for Biofilm Studies, and a Case Study of Its Application to Corynebacterium striatum, an Emerging Pathogen. Front Cell Infect Microbiol 2021;11. https://doi.org/10.3389/FCIMB.2021.803774.

[16] Vyas HKN, Xia B, Mai-Prochnow A. Clinically relevant in vitro biofilm models: A need to mimic and recapitulate the host environment. Biofilm 2022;4:100069. https://doi.org/10.1016/J.BIOFLM.2022.100069.

[17] Roy S, Santra S, Das A, Dixith S, Sinha M, Ghatak S, et al. Staphylococcus aureus Biofilm Infection Compromises Wound Healing by Causing Deficiencies in Granulation Tissue Collagen. Ann Surg 2020;271:1174–85. https://doi.org/10.1097/SLA.0000000000003053.

[18] Choong FX, Huzell S, Rosenberg M, Eckert JA, Nagaraj M, Zhang T, et al. A semi high-throughput method for real-time monitoring of curli producing Salmonella biofilms on air-solid interfaces. Biofilm 2021;3:100060. https://doi.org/10.1016/J.BIOFLM.2021.100060.

[19] Kadam S, Madhusoodhanan V, Bandgar A, Kaushik KS. From Treatise to Test: Evaluating Traditional Remedies for Anti-Biofilm Potential. Front Pharmacol 2020;11:1669. https://doi.org/10.3389/FPHAR.2020.566334/BIBTEX.

[20] Kadam S, Madhusoodhanan V, Dhekane R, Bhide D, Ugale R, Tikhole U, et al. Milieu matters: An in vitro wound milieu to recapitulate key features of, and probe new insights into, mixed-species bacterial biofilms. Biofilm 2021;3:100047. https://doi.org/10.1016/J.BIOFLM.2021.100047.

[21] Deresinski S. Vancomycin in combination with other antibiotics for the treatment of serious methicillin-resistant staphylococcus aureus infections. Clinical Infectious Diseases 2009;49:1072–9. https://doi.org/10.1086/605572/2/49-7-1072-TBL001.GIF.

[22] Post V, Wahl P, Richards RG, Moriarty TF. Vancomycin displays time-dependent eradication of mature Staphylococcus aureus biofilms. J Orthop Res 2017;35:381–8. https://doi.org/10.1002/JOR.23291.

[23] Malone CL, Boles BR, Lauderdale KJ, Thoendel M, Kavanaugh JS, Horswill AR. Fluorescent reporters for Staphylococcus aureus. J Microbiol Methods 2009;77:251–60. https://doi.org/10.1016/J.MIMET.2009.02.011.

[24] Shepherd J, Douglas I, Rimmer S, Swanson L, MacNeil S. Development of Three-Dimensional Tissue-Engineered Models of Bacterial Infected Human Skin Wounds. https://HomeLiebertpubCom/Tec 2009;15:475–84. https://doi.org/10.1089/TEN.TEC.2008.0614.

[25] Kadam S, Vandana M, Kaushik KS. Reduced serum methods for contact-based coculture of human dermal fibroblasts and epidermal keratinocytes. Biotechniques 2020;69:347–55. https://doi.org/10.2144/BTN-2020-0112.

[26] Abramoff MD, Magalhães PJ, Ram SJ. Image processing with ImageJ. Biophotonics International 2004;11:36–42.

[27] Ridler TW, Calvard S. PICTURE THRESHOLDING USING AN ITERATIVE SLECTION METHOD. IEEE Trans Syst Man Cybern 1978;SMC-8:630–2. https://doi.org/10.1109/TSMC.1978.4310039.

[28] Microsoft Excel Spreadsheet Software | Microsoft 365 n.d. https://www.microsoft.com/en-us/microsoft-365/excel (accessed February 2, 2023).

[29] Idrees M, Sawant S, Karodia N, Rahman A. Staphylococcus aureus Biofilm: Morphology, Genetics, Pathogenesis and Treatment Strategies. Int J Environ Res Public Health 2021;18. https://doi.org/10.3390/IJERPH18147602.

[30] Kim SJ, Chang J, Rimal B, Yang H, Schaefer J. Surface proteins and the formation of biofilms by Staphylococcus aureus. Biochimica et Biophysica Acta (BBA) - Biomembranes 2018;1860:749–56. https://doi.org/10.1016/J.BBAMEM.2017.12.003.

[31] Plikus M v., Wang X, Sinha S, Forte E, Thompson SM, Herzog EL, et al. Fibroblasts: Origins, definitions, and functions in health and disease. Cell 2021;184:3852–72. https://doi.org/10.1016/J.CELL.2021.06.024.

[32] El-Serafi AT, El-Serafi I, Steinvall I, Sjöberg F, Elmasry M. A Systematic Review of Keratinocyte Secretions: A Regenerative Perspective. Int J Mol Sci 2022;23. https://doi.org/10.3390/IJMS23147934.

[33] Klingstedt T, Åslund A, Simon RA, Johansson LBG, Mason JJ, Nyström S, et al. Synthesis of a library of oligothiophenes and their utilization as fluorescent ligands for spectral assignment of protein aggregates. Org Biomol Chem 2011;9:8356–70. https://doi.org/10.1039/C1OB05637A.

[34] Klingstedt T, Shirani H, Åslund KOA, Cairns NJ, Sigurdson CJ, Goedert M, et al. The structural basis for optimal performance of oligothiophene-based fluorescent amyloid ligands: conformational flexibility is essential for spectral assignment of a diversity of protein aggregates. Chemistry 2013;19:10179–92. https://doi.org/10.1002/CHEM.201301463.

